# A deep learning framework for quantitative analysis of actin microridges

**DOI:** 10.1101/2021.11.12.468460

**Authors:** Rajasekaran Bhavna, Mahendra Sonawane

## Abstract

Microridges are evolutionarily conserved actin-rich protrusions present on the apical surface of the squamous epithelial cells. In zebrafish epidermal cells, microridges form self-evolving patterns due to the underlying actomyosin network dynamics. However, their morphological and dynamic characteristics have remained poorly understood owing to lack of automated segmentation methods. We achieved ~97% pixel-level accuracy with the deep learning microridge segmentation strategy enabling quantitative insights into their bio-physical-mechanical characteristics. From the segmented images, we estimated an effective microridge persistence length as ~0.61μm. We discovered the presence of mechanical fluctuations and found relatively greater stresses stored within patterns of yolk than flank, indicating distinct regulation of their actomyosin networks. Furthermore, spontaneous formations and positional fluctuations of actin clusters within microridge influenced pattern rearrangements over short length/time-scales. Our framework allows large-scale spatiotemporal analysis of microridges during epithelial development and probing of their responses to chemical and genetic perturbations to unravel the underlying patterning mechanisms.

## INTRODUCTION

The apical surface of epithelial cells exhibits specialized actin-rich features. These include microvilli observed on intestinal epithelial cells that are required for absorptive function, and stereocilia present in the inner ear for mechanosensing (Blanchoin 2014, Apodaca 2018). Microridges are another class of actin-based protrusions found on various non-cornified squamous epithelia (Bereiter-Hahn 1979, Fishelson 1984, Uehara 1991, Depasquale 2018). They form laterally long labyrinthine patterns on the apical domain of peridermal or outer epidermal cell surfaces in the zebrafish embryos. Ultrastructural analyses have demonstrated that microridges are comprised of actin filament networks (Bereiter-Hahn 1979, Uehara 1991, Pinto 2019). Consistently, the function of Arp2/3 complex is essential for their formation and maintenance (Lam 2015, Pinto 2019, van Loon 2020). Additionally, actin regulators such as cortactin, VASP (Lam 2015), Wasl, Cofilin, Eplin, Filamin (Pinto 2019) and Non-muscle myosin-II (NMII) (Pinto 2019, van Loon 2020) as well as keratin cytoskeleton and Plakin cytolinkers (Inaba 2020) localize to the microridges. Microridges remain dynamic and actin is actively treadmilling within them (Lam 2015, Raman 2016, Sharma 2005). Besides, cell polarity proteins such as aPKC and Lgl - regulators of apical and basolateral domain identity, respectively - control the elongation of microridges (Raman 2016, Magre 2019). These studies provide collective evidence that microridges are organized by F-actin, NMII and regulated possibly by other actin binding proteins (ABPs) and cytoskeletal interactions. However, their precise interactions and mechanistic control during formation and maintenance remains elusive. As per the current understanding, F-actin punctae or pegs are distributed on the peridermal cells and apical constriction provide the necessary force for the neighboring pegs to coalesce into microridge (van Loon 2020), which gradually evolve into labyrinthine patterns, under the influence of Myosin-II activity (Pinto 2019, van Loon 2020, Raman 2016, van Loon 2021).

In reconstitution experiments, actin, NMII and their associated motor proteins can be organized into various large-scale patterns (Kruse 2004, Backouche 2006). The concentrations, kinetic parameters of ATP and density of actin related proteins contribute to their collective behaviour. Stable, stationary structures can form by self-assembly of actin and related proteins near their thermodynamic equilibrium. In contrast, in an active self-organizing system, different mechanisms can arise, in which the assembled structures reach an active steady state without reaching the thermodynamic equilibrium, as energy is continuously consumed and dissipated (Kruse 2004, Backouche 2006). In vivo, microridges remain in a non-equilibrium steady state by continuously reorganising their patterns constituted by an active network of F-actin, NMII and related proteins. One mode of understanding the mechanism of microridge formation and maintenance is to gain insight into their physical properties that are both reflective of and contribute towards the process of self-organisation. Spatiotemporal fluorescence imaging, segmentation and tracking form powerful approaches to gain quantitative insight into biophysical properties including morphological and dynamic characteristics.

Quantitative descriptions of dynamic processes require high quality image data followed by appropriate analysis methods (Bothma 2014, Castro-González 2014, Bhavna 2020). A major challenge for biologically relevant parameter extraction is image segmentation that correctly identifies pixels within the images. Depending upon image-content and morphological features such as cell membrane, nuclei or actin filaments, their size, shape, density and data dimension, rigorous image-based techniques are tailored to accurately address specific tasks and involve many carefully fine-tuned control parameters (Mosaliganti 2012, Bhavna 2016, Jacquemet 2017, Blin 2019). Often segmentation errors at small spatial scales can yield downstream errors leading to noisy results (Caicedo 2019). One way to circumvent this problem is by training machine-learning models on feature vectors extracted from annotated ground truth data representing all possible variabilities (Lucchi 2012, Berg 2019, Driscoll 2019, LeCun 2015). Another subset of machine learning algorithms are the deep learning methods that utilize the general principles of learning and in this process learn data representations with multiple levels of abstraction to discover intricate patterns required for detection or classification. They have remarkably outperformed feature-extraction based algorithms and surpassed human performance for hard problems (LeCun 2015). Specifically, convolutional neural networks (CNN) are designed to process multidimensional data arrays including multi-channel or temporal data sequences (LeCun 2015, Moen 2019). They have been applied with success for biological image analysis for cell types classification (Pliner 2019), protein subcellular localization (Kraus 2017), image restoration (Weigert 2018) and cell segmentation (van Valen 2016, Falk 2019, Lugagne 2020, Wang 2019).

We designed a fast and accurate large-scale CNN-based quantitative framework for analysis of microridge patterns. From the experimental data, we estimated the bending rigidity and population level length-scale parameter of microridges. Our flow analysis elucidated the time-dependent accumulation and dissipation of mechanical stresses within the underlying networks of microridge patterns. Our computational analysis of mobile high intensity actin clusters revealed their influence on localized pattern re-arrangements. Importantly, the framework allows a quantitative analysis of microridges, unravelling their mechanism of formation and maintenance at different developmental stages, response to perturbations and diverse genetic backgrounds.

## RESULTS

### Live imaging of zebrafish epidermis and image processing pipeline

The zebrafish epidermis is bi-layered by 48hpf with the apical surface of outer peridermal cells decorated with microridges. To facilitate high quality long-term live imaging of developing microridges, we designed and developed a custom mounting device by measuring zebrafish embryo dimensions at 48hpf (Methods). Microscopy parameters were optimized for achieving high spatiotemporal resolution of the epidermis from head, yolk and flank regions of the embryo (Fig 1a). Temporal changes in periderm cell height were sensitive to tissue regions and embryonic development. Therefore, optimal filtering parameters were set for each microscopy movie to encompass peridermal cells and exclude basal epidermal cells from the segmentation analysis (Fig 1b1, Supplementary Fig S1).

**Fig 1.**
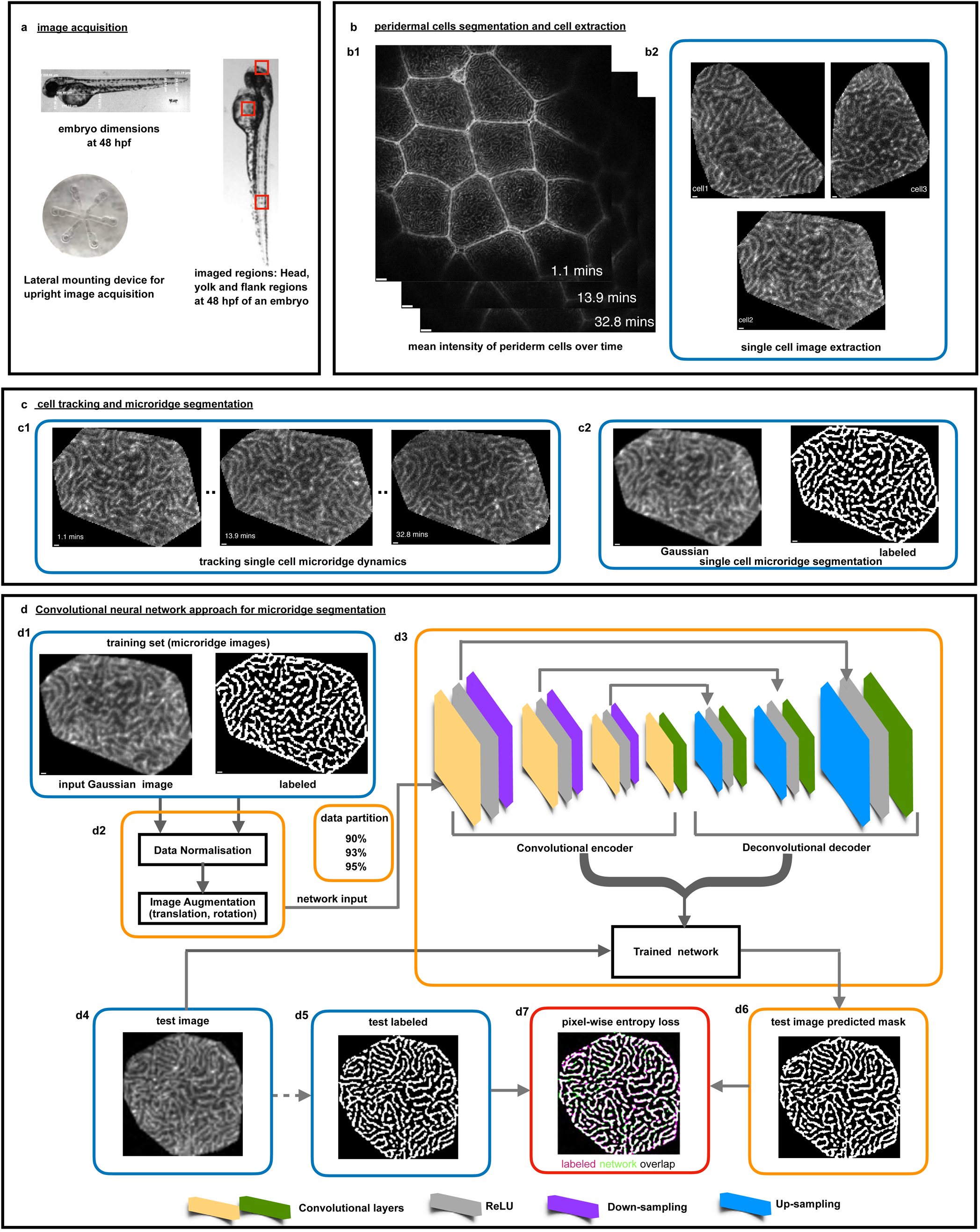
Live imaging, image processing pipeline for a neural network approach for microridge segmentation. **a**. Image acquisition. Zebrafish embryo dimensions were measured at 48hpf and a custom-built embryo mounting device was designed for live imaging of one lateral side of head, yolk, and flank regions of the embryo. **b**. Periderm cell segmentation and single cell extraction. **b1**. Mean intensity of the filtered periderm cell slices at all time points. **b2**. Membrane segmentation steps leads to demarcated cell boundaries and cell extraction. **c**. Cell tracking and microridge segmentation. **c1**. Nearest centroid distance-based cell tracking allowed following each cell’s microridge dynamics. **c2. A** fully-automated custom-built microridge segmentation algorithm formed the labelled set for the deep learning segmentation strategy (microridge image processing pipeline detailed in Fig S1, Supplementary Methods). **d**. Convolutional neural network approach for microridge segmentation. **d1**. Training set consisted of pairs of extracted cell pattern and their binarized images, described in b.-c. **d2**. Prior to training, the data normalization and data augmentation steps were implemented. Data was randomly partitioned into 90%, 93% and 95% of the total set and various combinations of hyperparameters are trialled in the training process. **d3**. The convolutional encoder-decoder architecture consisting of convolutional encoder and decoder layers (yellow and green), ReLU layers (grey), down-sampling (purple) and up-sampling layers (blue) yielded a trained network for each set of hyperparameters. **d4-5**. The network accuracy was assessed on the remaining test dataset (10%, 7% and 5% respectively) by pixel-wise comparison of network predicted output and labelled images. **d6**. Trained network predictions on test data. **d7**. Pixel-wise entropy loss was computed by comparing labelled images and network predicted images for assessing the network performance.

Custom algorithms were written for periderm cell-membrane segmentation followed by cell centroid-distance based tracking frame-by-frame. The cell segmentation steps excluded cells with incomplete edges (membranes) due to either low contrast or non-uniform z-fluctuations. Each cell mask demarcated a cell boundary (Supplementary Fig S1) and was used to extract raw single cells patterned with microridges (Fig 1b2). Cell-tracking information was used for dynamic pattern analysis. We designed an automated microridge segmentation pipeline that formed the labelled training set for CNN approach. The patterned-cell extraction step provided a large training dataset for CNN-based microridge segmentation (Figs 1c1-c2). Mathematical details of cell segmentation, tracking and microridge segmentation are described in Methods, Supplementary Methods and illustrated in Supplementary Fig S1.

### CNN based microridge segmentation: Tuning the training and performance evaluation

A step-wise description of the CNN microridge segmentation workflow is provided below (detailed in Methods). Using the training set, we optimized the hyperparameters to achieve a trained network by implementing a U-net encoder-decoder neural network architecture that has already demonstrated its efficiency in the bio-medical segmentation field (van Valen 2016, Falk 2019). The scheme in Fig 1d illustrates the deep learning approach for microridge segmentation.

#### Training set

The training set consisted of image pairs of raw cell patterns and their corresponding labelled image annotations using the automated microridge segmentation pipeline. The number of pixels that amounted to foreground and background were variable across different patterns. The dataset formed an excellent repository for training the network as it offered important characteristic features owed to variations in patterns. These in turn served to determine the optimal set of hyperparameters to achieve high pixel-wise segmentation. (Fig 1d1).

#### Optimizing hyperparameters

We applied median pixel image normalization to balance the weight of foreground and background pixels during training to solve the binary pixel-classification problem. Having produced microscopy data within the laboratory, a data augmentation step was implemented (Methods, Fig 1d2) to ensure that sufficient information was provided to the network for training. The network performance was sensitive to hyper-parameters. After initial testing of several combinations of hyperparameters, we fixed the image size to 256^2^ (pixels), which scales to the receptive field size and encoder depth of 6 and adjusted the learning rate to 10^−4^. Smaller receptive fields less than 6 led to higher numbers of falsely classified pixels, whereas optimal performance was obtained at the cost of longer training time. We varied the mini-batch size (MBS) that indicates the subset of data from training set that is used at each training iteration and maximum epochs (ME), which is the number of iterations through the entire training dataset during the training.

#### Network setup for training

We performed numerical tests by varying i) the fraction of the total dataset during training (95%, 93%, 90%) and ii) MBS and ME for each data proportion (Fig 1d3). The trained network performance was evaluated by measuring the accuracy on the remaining unlabelled test images (5%, 7%, 10%) whose image pixels were assigned independently by the microridge segmentation pipeline (Figs 1d4-d5). The network training time varied from 12-20 hours depending upon the combination of hyperparameters (e.g., training time increased with larger ME) on a GPU enabled high performance cluster.

#### Performance evaluation

The trained networks predicted segmentation on test images (Fig 1d6). For each set of hyperparameters, we computed the pixel-wise entropy loss to evaluate the segmentation accuracy (Csurka 2013) of microridges by pixel-wise comparison of labelled image with network predicted output using the weighted intersection over union (Weighted IOU) that weighted the number of pixels for each class (Fig 1d7). For most cases, an accuracy of about ~90% was achievable. The numerical tests indicated higher segmentation accuracy for smaller MBS and larger ME rather than otherwise (Fig 2a), suggesting that a repeated learning process was better than giving a greater number of images in one iteration for such a segmentation task. We selected the network trained with 93% data proportion, MBS=6 and ME=800 that yielded a ‘weighted mean IOU’ of 97.3% (Fig 2a) for quantitative pattern analyses.

**Fig 2.**
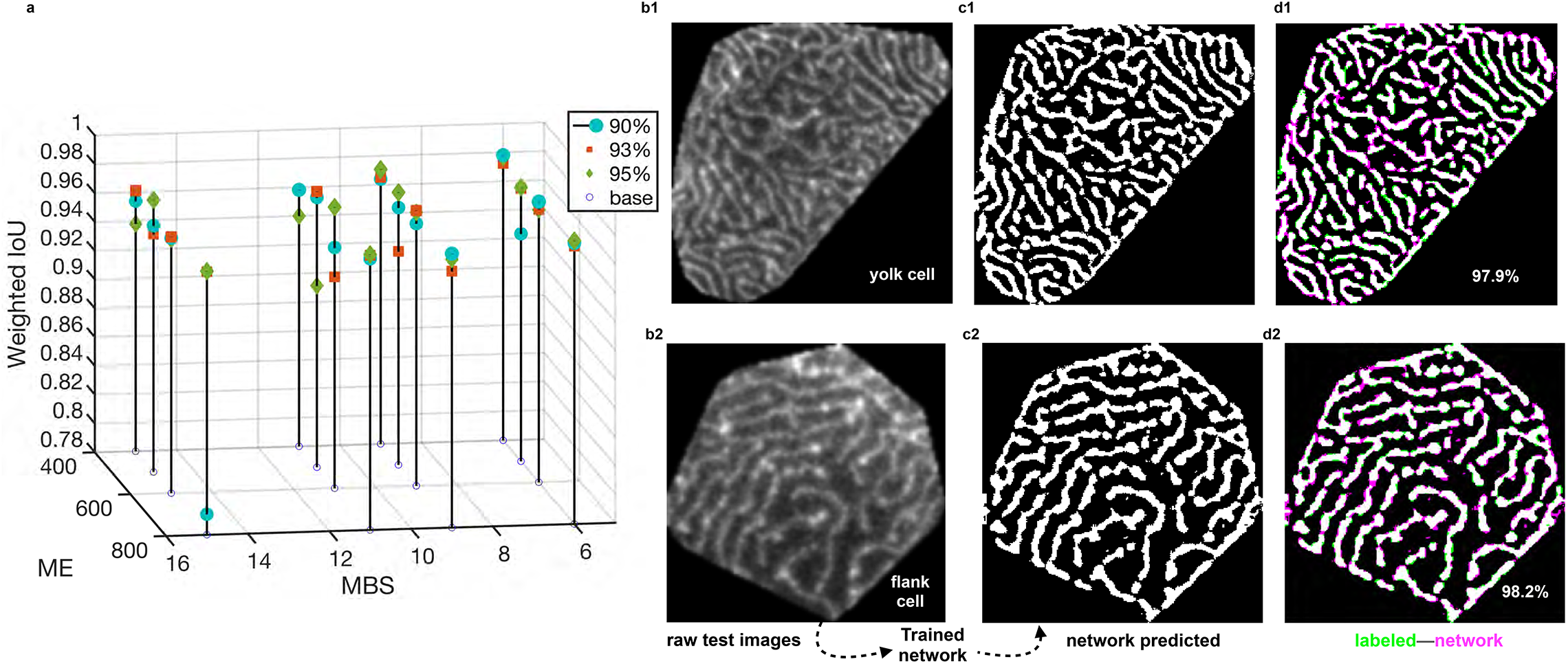
Trained network selection based on predicted accuracy versus network hyperparameters and visual inspection of pixel-wise segmentation. **a**. 3D stem plot shows how the mini-batch size (MBS) and maximum epoch (ME) affect the network performance, measured by the weighted IOU (Average intersection over the union of all classes, weighted by the number of pixels in the class). For each proportion of the training set, the MBS and ME values were varied as (6, 9, 11, 15) and (400, 500, 600, 800) to yield 16 combinations of these hyperparameters. Typically, better accuracy is achievable for smaller MBS and a larger ME, for which our tested combinations of hyperparameters showed performances above ~90% weighted IOU. **b1-b2**. An exemplary single yolk and flank pattern respectively, fed to the selected trained network. **c1-c2**. Trained network segmented outputs for MBS= 6 and ME=800 using the 93% training dataset (1397 randomly chosen microridge patterns). **d1-d2**. Pixel-wise overlap between images labelled using the conventional microridge segmentation pipeline and network segmented images, shown in green pixels and magenta pixels respectively; common regions in white (microridges) or black (background). Performance measure given by weighted IOU for each cell pattern.

A visual comparison of yolk and flank cell patterns with the corresponding network segmented outputs indicated reasonable performance (Figs 2b1-c1, 2b2-c2, 2d1-d2). Supplementary Movies 1 and 2 show the time evolution of microridge patterns of raw and network segmented images for a yolk and flank cell, respectively. For all network segmented image pixels, we obtained their physical sizes prior to quantitative analyses (Supplementary Methods). Below, we describe the static parameters of microridges, followed by examination of their steady state dynamics.

### Estimation of bending rigidity of microridges *in vivo*

Deciphering the chemo-mechanical properties of microridges is pertinent towards understanding emergence and maintenance of their patterns. We estimated their effective persistence length (*L_p_*) or the characteristic length scale at which a microridge bends, to assess their inherent mechanical stiffness. The mechanical responses such as bending, stretching or compression of the in vivo microridges is likely to be a consequence of active fluctuations in actomyosin network forces, in addition to their dynamic response to thermal fluctuations. Previously, such measurements have been established in *in vitro* single actin filaments and microtubules, purified extracts of DNA polymers based on their thermal response (Gittes 1993, Rivetti 1996), and microtubules within cells as a consequence of both thermal response and cytoskeleton elements (Brangwynne 2007).

We defined an effective *L_p_* based on the overall curvature distribution of microridges (Methods). For each 2D sub-image of a skeletonised microridge branch, (Fig 3a, highlighted within a magenta box), we obtained discrete x–y pixel coordinates traced along the skeleton contour length (*L_xy_*) (Fig 3b inset) that was fitted with a cubic spline to obtain a smooth microridge contour (Eqs. 1–3, Fig 3b). For each curved segment, we computed the spacing (Δ*s_k_*) between adjacent points on the skeleton and the tangent orientation angle (*θ_k_*) (Fig 3c), which together allowed the evaluation of the corresponding local curvature (*κ*) (Fig 3d) along the contour length (Eqs. 4–8). We rescaled the curvature (*κ*) by the square root of the local segment length to *κ_s_* (Fig 3e, Eq. 9) to extract the effective persistence length (*L_p_*). We fitted a Gaussian into the distribution 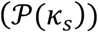 (Eq. 10, Methods, Fig 3f), whose width defines an effective *L_p_* estimated as ~0.61μm. We determined the effective flexural rigidity (Eq. 11) or bending rigidity of microridges as 2.54×10^−15^ Nm^2^. The internal forces generated by actomyosin networks within the microridges govern their response under active load, such as stretching, compression or even buckling. For an isolated microridge of length *L~*1μm, with the estimated *L_p_*, under active load, the critical force (*f_c~_ π^2^k_B_TL_p_/L^2^*) would be about 0.025 pN, above which the microridges would readily buckle. This critical force is less than for pure actin filaments that have *L_p_* ≈17 μm (Gittes 1993) and thus *f_c_* of around 0.69 pN. Our analysis is based on the mesoscopic properties of the network within the microridges. A bottom-up approach, such as building in vitro (or reconstitution) models, would require mimicking the meshwork properties of microridges. Hence, an estimate of *L_p_* describing the mechanical properties of the molecular network is fundamental to probing the role of constituent proteins and deciphering their mechanisms.

**Fig 3.**
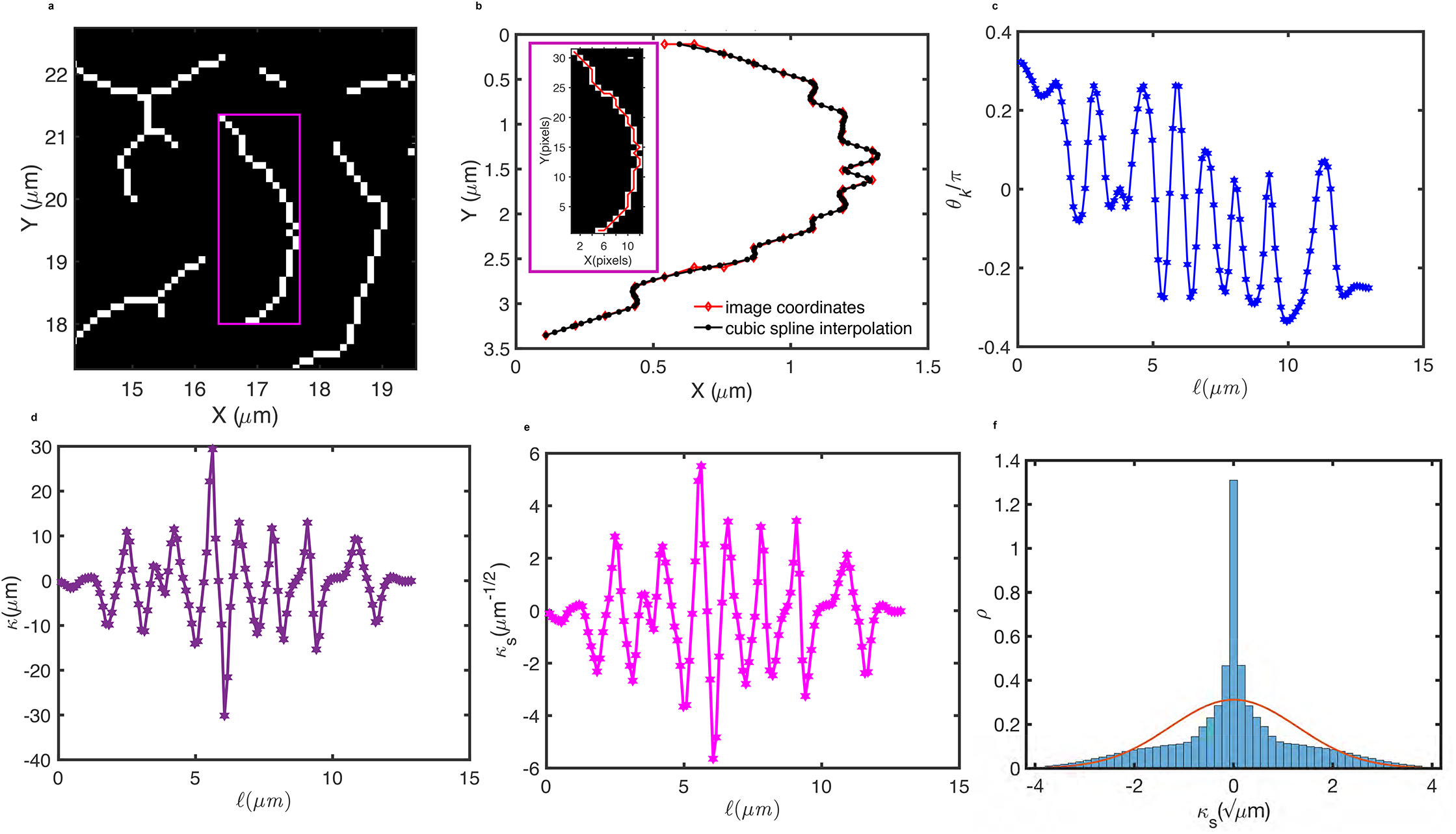
Estimation of persistence length for in vivo microridges. **a**. A magenta box demarcates the 2D sub-image of a skeletonised microridge branch for estimation of L_p_. **b**. Image coordinates shown in red with the best fitting ‘cubic smoothing spline’ fit and the inset shows the discrete x–y coordinates obtained along the traced microridge (red line) of skeletonised contour. **c**. Tangent angle (*θ_k_*) along the length (*ℓ*) of the microridge. **d**. The local curvature (*κ*) of the microridge along its length (*ℓ*) was calculated from the tangent angle. **e**. Rescaled *κ_s_* along the length (*ℓ*) of the microridge. **f**. Distribution of *κ_s_* of microridges from 593 cells (300 yolk cells and 293 flank cells) fitted to a Gaussian distribution (red line trace), whose variance gives an estimate of the effective persistence length (*L_p_*) as ~0.61μm.

### Population level microridge pattern length-scales

The emergent pattern of microridges is a signature of their underlying molecular determinants. Hence, the concentration, diffusion and degradation rates of actin and interacting proteins can affect the spacing, density and thereby the pattern length-scale and pattern span characterized by the pattern wavelength. We estimated the wavelength (*λ*) in the Fourier domain (Eq. 12, Methods) for cell patterns from both yolk and flank regions, representative patterns are shown in Figs 4a-b respectively. The median values for yolk and flank cell patterns were found to be 0.66μm and 0.60μm respectively (Fig 4c). The spatial arrangements of cell patterns from yolk and flank regions were visually discernible with yolk cell patterns being more crowded, with several short-length microridges within centre and longer microridges along the cell periphery, whereas flank cell patterns typically comprised of less crowded but longer microridges. We determined microridge mean branch lengths (<*B_l_*>) for each patterned cell (Methods) and found relatively less variance within the yolk population than in patterns from flank regions as indicated by the boxplots in Fig 4d.

**Fig 4.**
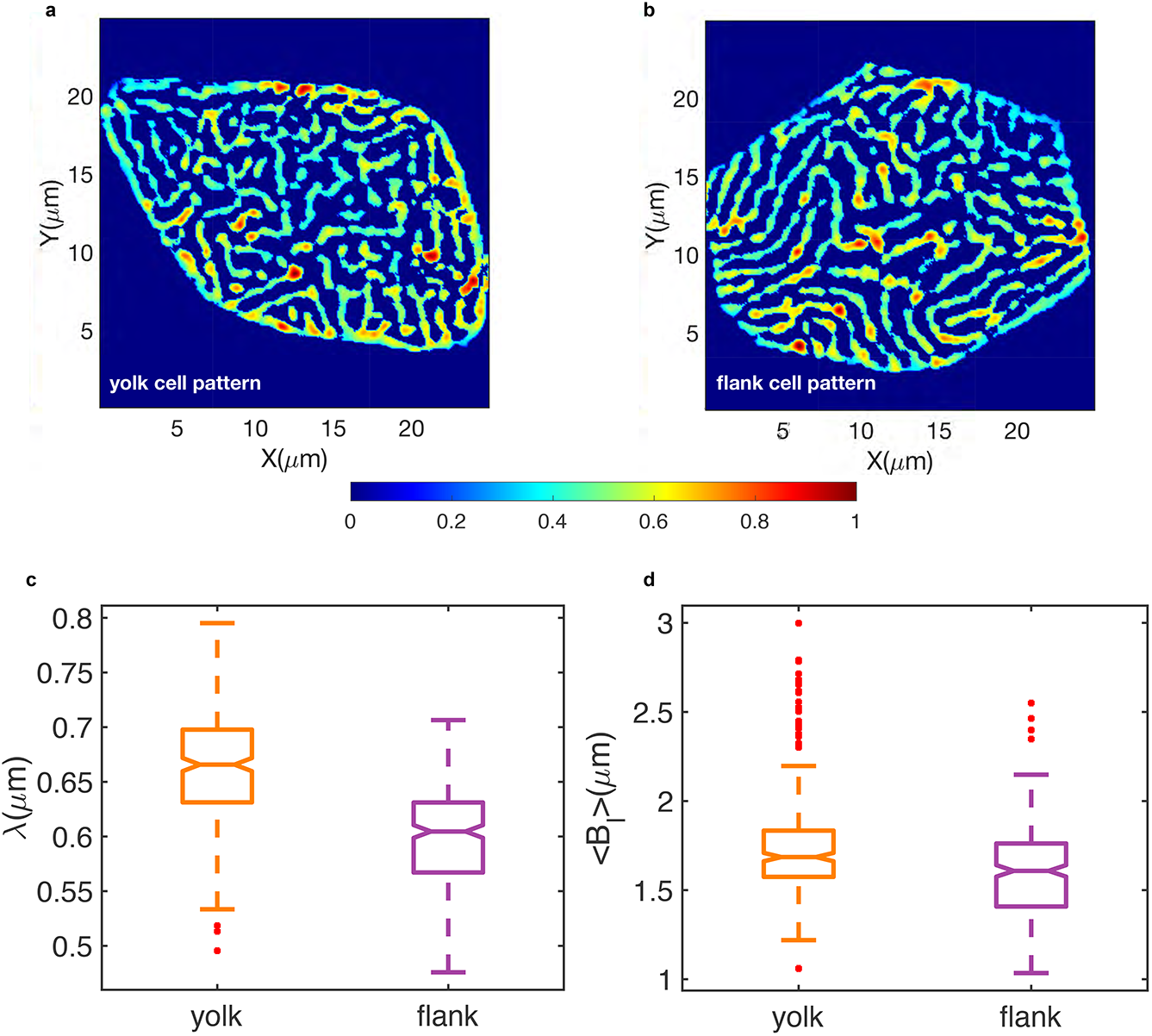
Population level comparison of cell patterns from yolk versus flank regions Example of a network segmented. **a**. yolk pattern **b**. flank cell pattern, both shown in false colour representing image intensities scaled between 0-1, indicated by the colorbar. **c**. Box plot of pattern wavelength (*λ*) parameter with estimated medians of 0.66μm and 0.60μm measured from network segmented binary cell images of yolk and flank regions computed over 300 yolk and 293 flank cells respectively. **d**. Box plots of yolk and flank cell population mean microridge branch lengths (<*B_l_*>) per cell respectively quantified from their skeletonised microridge branches.

### Microridge flow fields reveal complex interactions of internal active forces generated by actomyosin networks

Actin networks driven out of equilibrium by force generation through NMII activity can lead to mechanochemical pattern formation (Howard 2011, Prost 2015, Ben Isaac 2013). The interplay between active force generation and force dissipation in a visco-elastic environment governs the dynamic properties of such a system and the active stresses could regulate steady flow patterns (Howard 2011, Prost 2015). We performed a spatiotemporal flow analysis of the evolving microridge patterns to gauge the complex bulk dynamics and gain insights into the underlying network remodeling process near 48hpf. In order to estimate the degree of mechanical stresses, we evaluated the local deformations of the actin networks by characterizing the pattern velocity flow fields and the corresponding local strain rates. We used an optic flow method, assuming image intensity flow corresponded to material flow. In order to demonstrate the flow analysis, we selected a square region within a yolk cell center patterned with microridges (Supplementary Movie 3a, Fig 5a). The microridge configuration at two consecutive time points, t_1_=0 and t_2_=0.6 mins are shown in Figs 5b-c. The microridge pattern at t_1_ superimposed by velocity vectors (magenta arrows) indicating flows between consecutive time points from t_1_ to t_2_ is illustrated in Fig 5d. Supplementary Movie 3b panel-1 shows the complete time evolution of the flow velocity (Eq. 13) for the yolk cell pattern shown here. The regions with longer magenta arrows (Fig 5d) move farther, signifying active forces arising from molecular interactions driving the dynamic bulk events. These events were: i) elongations caused by pulling in the outward directions along its lengths, or steered by fragmentation at the ends of a microridge, either events leading to change in length of a single microridge, ii) merging or splitting (similar to fusion or fission (van Loon 2021)) involving several microridges. To examine these events, we computed the velocity field divergence 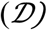, (Eq. 14) shown in Fig 5e and Supplementary Movie 3b, panel-2. High positive 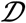 (red) represented regions with the loss of underlying components within the microridges, hence local flow moving out from a region (fragmentation or splitting events, considered together as shrinkage event), while low negative (blue) represented regions with gain of microridge pixels or local flow coming into a region (elongation or merging events, considered together as growth event) between consecutive timepoints. To focus exclusively on growth or shrinkage events, we determined the 2D spatial coordinates with large magnitude of 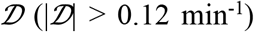 and extracted the velocity magnitude in these regions. We then estimated the growth velocity (*υ_growth_*) and shrinkage velocity (*v_shrinkage_*) (Eq. 13–14) by averaging them over the entire cell pattern and all frames. We found (2.33±1.39)×10^−2^ μm min^−1^ and (2.34±1.39)×10^−2^ μm min^−1^ for the yolk pattern frames, and (2.04±1.34)×10^−2^ μm min^−1^ and (2.05±1.35)×10^−2^ μm min^−1^ for the flank pattern time series, respectively (Supplementary Movies 1-2).

**Fig 5.**
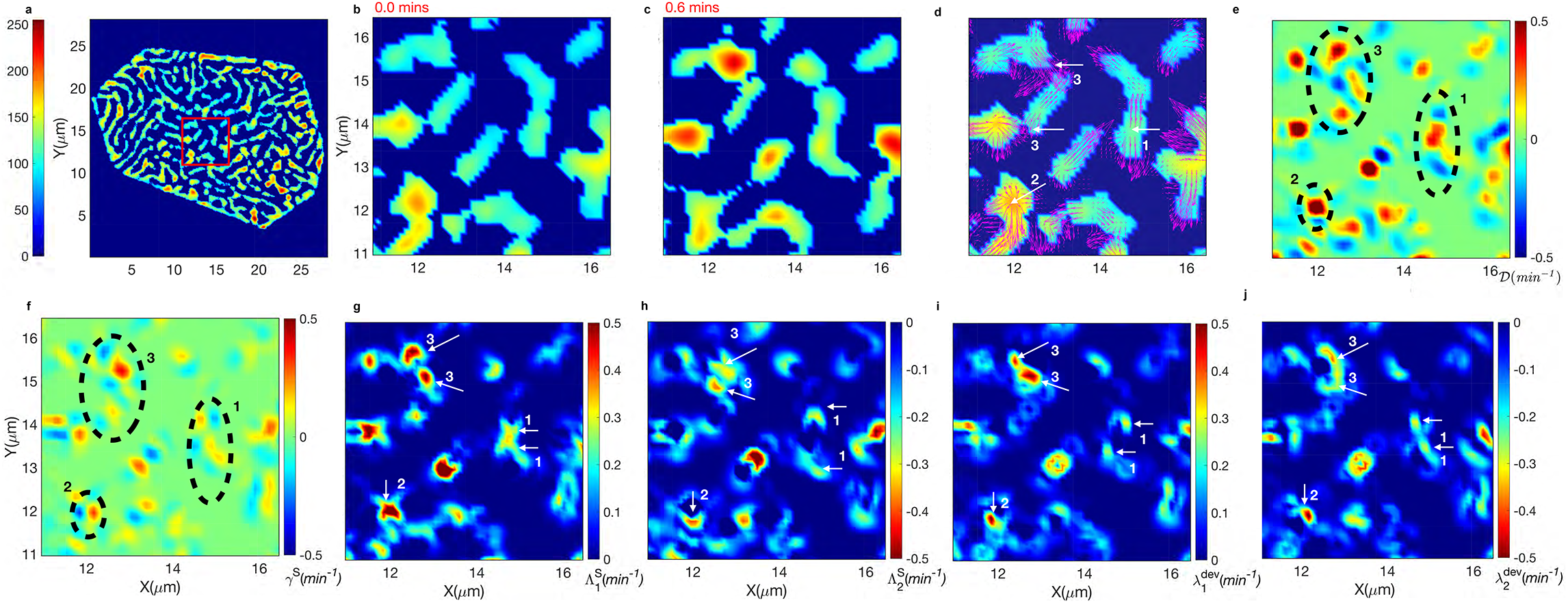
Microridge velocity flow field analysis reveals localized strain rate components within actin-myosin network for generating active flow patterns. **a**. A representative network segmented cell with image intensities shown in false color. A preselected squared center region in red is used to demonstrate the flow analysis. **b-c**. Microridge pattern at t_1_=0 mins and t_1_=0.6 mins respectively. **d**. Velocity vector field as magenta arrows, indicating the flows from t_1_ to t_2_, shown overlaid on the pattern at t_1_. Higher velocities (larger arrows) coincide with either elongation (from within) or fragmentation (at the ends) of a microridge, or merging or splitting of multiple microridges (shrinkage or growth events). White arrows (1-3) indicate pre-selected regions for detailed description of divergence and strain rates within the microridges shown in e, f and g. **e**. Divergence, 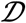, shown within the squared region of (a), reaching both large positive and large negative (colorbar), indicating regions with fragmentation or splitting and elongation or merging respectively. Encircled regions 1-3 indicated regions of elongation, outflows and inflows respectively. **f**. Shear strain rate (*γ^S^*) shown within the same regions. **g**. Tensile strains 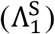 indicated by arrows 1-3, transiently developed adjacent to regions of elongation or next to outflow or inflow regions. **h**. Localized compressive strains 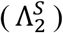 adjacent to tensile strains indicated by arrows 1-3 respectively. **i**. Deviatoric elongation strains without change in local area 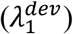 indicated by arrows 1-3 respectively. **j**. Strains representing deviatoric shrinkage without change in local area 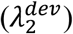 indicated by arrows 1-3 respectively. Here, *S* and *dev* denote strain rate tensor components and deviatoric strain tensor components respectively.

The flow analysis provided mesoscopic pointers towards the internal stresses within the meshwork of actin, NMII and related proteins involved in the complex dynamic flows of actin microridges. In principle, the actin cytoskeleton in the presence of NMII consists of network filaments that could bear both tensile and compressive forces. To examine the local material sources (large positive 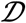) and adjacent sink (large negative 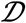) regions, we quantified the strain rate tensor (*S*) components (Eq. 15–17): shear (γ^S^), tensile 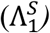 and compressive 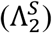 strains, respectively (Figs 5f-h). We additionally computed the deviatoric tensor *S^dev^* (Eq. 18–21) that estimated pure elongation 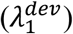 and pure shrinkage 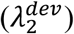 of microridges excluding local area changes (Figs 5i-j).

We compared the same regions within Figs 5d-j, marked by dashed circles or arrows 1-3. The arrow-1 in Fig 5d shows microridge elongation that necessitates two opposite local velocity fields generating two localized +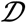 regions flanked by two adjacently located outer −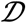 regions accompanied by γ^S^ shear strain rates (area encircled by the dashed line-1 in Figs 5e-f). The complex internal strain fields exhibited two local tensile strain regions 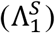 and adjacent compression regions 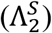, additionally steered by deviatoric elongation 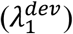 and deviatoric shrinkage 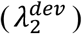 respectively, indicated by their colorbars (arrows-1 in Figs 5g-j). The localized strains transiently build-up and dissipate immediately in the neighboring region. Next, we examined arrow-2 (indicating a merging event of two microridges) in Fig 5d with outflow velocities in all directions from a single region and hence positive 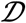 (dashed-circled region-2 in Fig 5e), leading to localized tensile strains (arrow-2, Fig 5g) and neighbourhood compressive strains (arrow-2, Fig 5h). The deviatoric shear components are shown by arrow-2, Fig 5i-j. In the dashed-circled region-3 and arrows-3 in Figs 5d-j, we observed multiple directed flows resulting in complex strains in very nearby regions. The time evolution of all flow parameters for the yolk pattern is shown in Supplementary Movies 3b. The microridges exhibited resistive viscoelastic forces alternating between rapid extensions (resistance to outflow) and compressions (resistance to inflow) hence exhibiting temporal periodic fluctuations. The area-deviatoric decomposition of the S-tensor indicated that deviatoric shear strains within the networks including sliding-movements and localized transient area changing events both contribute to microridge flows.

We computed the overall principal strains 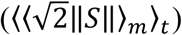 for the cell patterns, temporally averaged, and found them to be (11.15±2.16)×10^−2^ min^−1^ and (9.77±1.64)×10^−2^ min^−1^ for the yolk and flank cell patterns (Supplementary Movies 1-2), respectively. The comparative pattern strain rates and their mean velocities for the analyzed patterns are listed in Supplementary Table S1. The presence of temporal periodic fluctuations of the mechanical parameters within microridge patterns are shown in Supplementary Fig S2. Our results showed relatively greater strain fields within the actomyosin networks of microridge patterns of yolk than flank cells, suggesting that the stress distributions could be distinctly regulated in different regions of the same embryo. The time evolution of flow parameters within a flank cell pattern are shown in Supplementary Movies 4a-b.

### Intensity profile within each microridge remains dynamic

Interestingly, we observed high intensity spots traversing along the microridge lengths, indicating clustered actin speckles exhibiting positional fluctuations. We performed microridge tracking and computationally analyzed the intensity profile along the microridge lengths (Eq. 22). An exemplary four-time frames of a single microridge are shown (Figs 6a-d). Their intensity profile indicated movement of high peaks due to a mobile actin protein cluster within the microridge (Fig 6e). The presence of high intensity spots is also confirmed by a high-resolution STED microscope (Fig 6f). The intensity profiles along another microridge are also shown in Supplementary Movie 5. Plausibly, high intensity clusters become unstable and hence fluctuate in position. This may result in transient and localized spontaneous positive microridge curvatures orthogonal to the cell surfaces. These spontaneous positive curvatures could lead to instabilities and height fluctuations (Gov 2006). To analyze the fluctuating intensity spots, we computed the localized Gaussian curvature (*K*) considering microridge intensity profiles to be proportional to their heights (Eq. 23). Supplementary Movie 6 shows the intensity profile within a sub-region of a pattern and the computed localized curvature whose z-profile corresponds to intensity.

**Fig 6.**
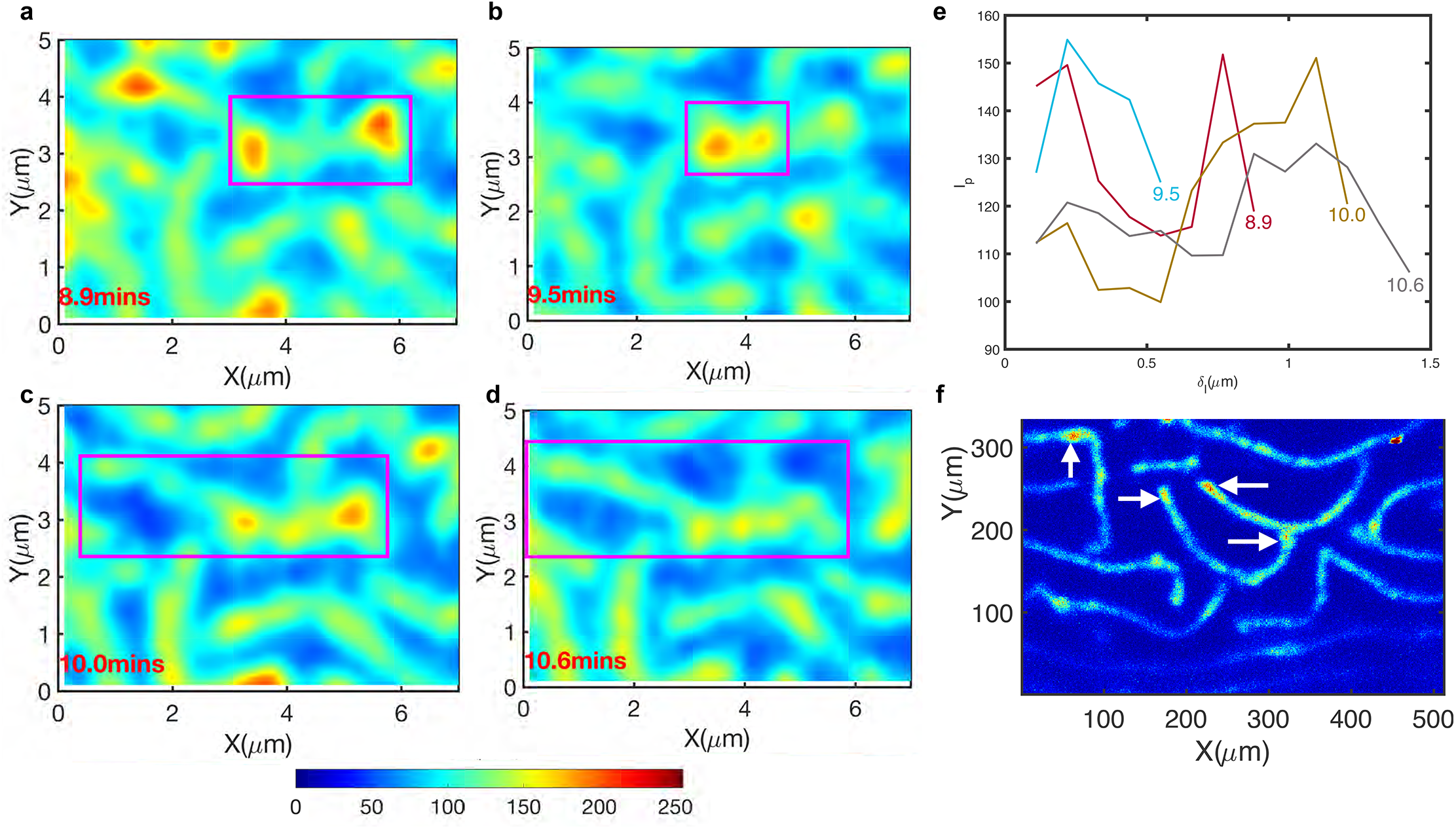
Actin clusters within the microridges exhibit positional fluctuations. **a.-d**. The raw image intensity (in false color) of a representative microridge indicated the presence of actin clusters traversing along the microridge lengths across time frames. **e**. Microridge tracking allowed extraction of one-dimensional intensity profile along the same microridge at each timepoint. Each line color indicates the intensity profile, labelled by the time in minutes, showing the intensity fluctuations within the microridges. High intensity peaks oscillate in position along microridge lengths. **f**. A high-resolution STED image at resolution of 0.022μm in x and y and z-resolution of 0.22μm shows the clustered intensity spots within the microridges labeled with Utr-gfp.

We determined the number of 2D-pixel coordinates of high positive and low negative curvatures (|*K*|>5) and found localized high *+K>5* constituted ~70% of these, on average per cell pattern at all timepoints. We then carried out time-evolution of 2D-pixel location coincidence analysis between ±*K* (intensity) and ±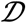 locations (significant flows). We found high frequency counts of *+K* locations at time point *t* overlapped with significant flow locations from timepoint *t* to *(t+1)* (Supplementary Fig S3). The positional fluctuations of *+K*-locations in the form of high intensity clusters continuously altered adjacent sink/source regions within a microridge and could influence localized pattern rearrangements in the form of shrinkage or growth events over short time scales. The short transient time between accumulation and dissipation of mechanical stresses could influence the formation and mobility of high intensity clusters at 48hpf to recruit or retain other ABPs that may influence the turnover and binding constants of subunits of actin. However, the precise molecular cause and the mechanism that drives this process remains to be determined.

## DISCUSSION

Image segmentation methods are central to transforming live-imaging experiments into quantitative information, which facilitates better understanding of the dynamics of processes of interest. Our cell segmentation, tracking and single cell extraction from microscopy images are generic methods adaptable for quantitative studies of other cellular processes. Recently, deep learning algorithms for cell segmentation have used manual annotations (van Valen 2016, Falk 2019) for obtaining labeled training datasets to mask cells. However, manual annotation can become increasingly laborious for annotating cell surface patterns and prone to errors. Hence, we built an automated algorithm to label microridges for the training set. Using the best performing trained network, we achieved microridge segmentation per cell in less than a minute. Improved microridge segmentation accuracy using CNN strategy consequently improved data quality required for in-depth quantitative analyses.

The effective persistence length (*L_p_*) indicated the nature of the cytoskeletal network underlying microridges in vivo. We estimated *L_p_* of *in vivo* microridges to be ~0.61μm, smaller than the microridge contour lengths. Under in vitro conditions, pure microtubules, actin filaments and DNA show *L_p_* of about 6mm, 10μm and 50nm and behave as rigid rods, semi-flexible polymers and very flexible polymers respectively (Gittes 1993, Rivetti 1996). When actin filaments interact with other associated proteins, their *L_p_* is considerably altered. In vitro studies on F-actin and Fascin–actin bundles showed significantly less thermal bending fluctuations in Fascin associated actin than in F-actin. The rigidity was sensitive to the ratio of F-actin and Fascin bundles, as the stiffness increased with the increasing concentration of actin bundling protein (Claessens 2006, Takatsuki 2014). On the other hand, cofilin-decorated actin filaments showed 5-fold lower *L_p_*, suggesting greater flexibility than native filaments (McCullough 2008). Further, actin filaments showed reduced *L_p_* in the presence of phalloidin or heavy meromyosin as compared to bare actin (Bengtsson 2016). These studies together indicated that interactions of actin with Myosin or ABPs significantly affected their flexibility and mechanical properties. The overall mechanical properties and the resulting force-related parameters would be modulated by the molecular components within the microridges resulting in lower *L_p_*. F-actin pegs shifted out of equilibrium possibly due to apical constriction (van Loon 2021) are driven into a steady state microridge patterning process. Our work suggests additional factors that could influence the evolution of microridge patterns. Our spatiotemporal analysis indicated that 2D planar flows driven by both extensile and compressive forces and steered by localized shear strain rates that are present within the microridges at 48hpf. Temporal periodic fluctuations of mechanical parameters could have implications in mechanochemical feedbacks involving Rho family GTPases (Martin 2014) known to drive contractions of the actomyosin cortex in other contexts. Further, we found greater strain fields within the microridge patterns on the periderm cells over the yolk than the flank regions that can be attributed to various physical and molecular differences present in the two regions. For example, junctional organization on yolk versus the flank does not develop in the same manner (Sonawane 2005), suggesting regional differences in signaling mechanisms. The yolk periderm cells are held at a greater height (from the basal epidermis) and reside over curved surfaces (attributed to overall tissue curvature) in contrast to flank cells. These together in turn could rework the underlying active localized stress distributions within the cortex, thereby distinctly remodelling the actomyosin networks and regulate microridge pattern scaling and dynamics. Our comparison of yolk versus flank pattern strain fields revealed that apart from generic features of mechanochemical patterning (van Loon 2020), the milieu in which cells reside could regulate the stresses within the actomyosin networks.

The low L_p_ indicated that the net force parameter of microridges is smaller in comparison with other classes of actin-based protrusions. If the protrusive forces of polymerizing actin are not large enough, then such a force could result in microridge growth in the lateral direction rather than causing significant increase in their heights. We observed and analyzed transient high intensity actin clusters in the form of spontaneous positive microridge curvatures that exhibited positional fluctuations along the lateral directions and traversed at a speed of upto ~2.8μm min^−1^ along microridges lengths. We anticipated that these unstable clusters influenced the flow events. Concomitantly, we found 2D-spatial locations of high intensity clusters coincided with locations of significant flows at consecutive timepoints, suggesting that dynamic clusters could sporadically lead to either growth or shrinkage event.

Although not determined experimentally, the collective interaction of actin regulators could lead to formation of spontaneous positive curvatures within microridges exhibiting instabilities and height fluctuations. Recent works have elucidated the role of NMII minifilaments in microridge remodelling by promoting fission and fusion events (van Loon 2021). Reconstitution studies have depicted a novel picture of NMII mini-filaments in organizing various dynamic patterns (Kruse 2004, Backouche 2006, Köhler 2011, Soares e Silva 2011). NMII activity could drive a network by a multistage coarsening process with NMII foci processively running over actin clusters resulting into a dynamic steady state (Soares e Silva 2011). This process could lead to actin filaments into disorganized condensates characterized by a broad actin cluster-size distribution (Köhler 2011). As a result, at certain concentrations, actin clusters subjected to NMII motors may themselves become highly mobile (Köhler 2011). It is likely that the interaction of NMII motors with F-actin and other actin regulators like Arp2/3 results in dynamic high intensity clusters within the microridges. The positional fluctuations of actin clusters could be a signature of the dynamic re-structuring of the relatively disorganized actomyosin network around 48hpf that dictates pattern rearrangements over short length/time-scales. As the pattern matures, microridges organise into a parallel fashion by 96hpf (van Loon 2021). Further investigations are required to link the molecular aspects of transient clusters driving actin flows during pattern dynamics.

To summarise, our CNN framework allows large-scale quantitative analyses to decipher the mechanisms of microridge pattern evolution and maintenance. We have identified some of the key aspects influencing microridge pattern dynamics. The persistence length of microridges indicated the range of their force parameters. Our comparative flow analysis elucidated patterns varyingly evolve on different regions of the embryo thereby bringing forth the significance of distinct regulation of mechanical stresses within the actomyosin networks. Furthermore, we discovered transient presence of high intensity clusters that influenced pattern flows near 48hpf. Further experimental and theoretical investigations are required to understand the complex interplay of actin, their associated proteins, the role of mechanics, cell morphology and the precise role of feedback signaling mechanisms in shaping the emergent pattern of microridges and their dynamics. Our CNN based segmentation framework will be immensely useful in the future in these efforts.

## METHODS

### Zebrafish strains

All zebrafish (*Danio rerio*) husbandry and experimental procedures were performed in Tübingen (Tü) strain. We used previously characterised zebrafish lines *Tg(actb1:GFP-utrCH)* provided by Behrndt and colleagues (Behrndt 2012). Zebrafish were raised and kept under standard laboratory conditions. The zebrafish maintenance and experimental protocols used in this study were approved by the institutional animal ethics committee.

### Mounting zebrafish embryos for live image acquisition

Zebrafish embryos were mounted for live imaging to record the dynamics of microridges on the periderm cells from 2-2.5dpf stages. An agarose-free flat mounting setup, custom fabricated at the TIFR workshop, was used for imaging the lateral side of the embryo. Live imaging was performed in E3 buffer with 0.02% ethyl-m-aminobenzoate methanesulphonate (Triacane). Live images of the microridges on the periderm cells within the yolk, head and flank regions of the embryo were obtained using a 40x dipping lens upright Zeiss confocal 880 microscope. The z-stack images from apical upto basal epidermis were continuously obtained at time intervals of around ~0.5 mins for each region during live image acquisition. The images were acquired at a preset room temperature with fluctuations between 25°C - 27°C.

### A custom-built image processing pipeline for microridge segmentation

A two-step segmentation and tracking approach was implemented. Firstly, we built a custom method for cell membrane segmentation followed by cell tracking. Single cells were isolated with their microridges using cell boundary information. Cells were tracked by using the nearest neighbor cell centroid distances for frame-by-frame cell association and assignment. Single cell isolation and subsequent tracking allowed to follow the same associated cell over all time frames and thereby also follow its intrinsic microridge dynamics (Supplementary Methods). For pixel-wise identification of microridge patterns, a second custom-built segmentation algorithm was designed with a series of image processing techniques to produce a segmentation mask of the microridges where each pixel was assigned a label for microridges and another label for background pixels (Supplementary Fig S1, Supplementary Methods). The labeled microridge were used for training a convolutional neural network (CNN) for segmentation. Manual segmentation was not considered for microridges, owing to their small size and manual errors, however the labeled data was visually verified with raw data. For the CNN segmentation approach, the entire dataset was provided during training (without cell tracking information). The cell tracking approach was implemented for the computation of microridge parameters during their steady state dynamic analysis.

### Training datasets for convolutional neural network microridge segmentation

The extracted raw microridge cell images formed the input for training the segmentation network. Corresponding annotated images of microridges consisting of foreground pixels (microridges) and the background pixels formed the labelled training set to determine the network parameters. We used a semantic segmentation approach, which converts the image segmentation problem into a pixel-level image classification problem. The task was reduced to finding a binary classifier for each image of the training set. Using this approach, we constructed a labelled training set with images of microridges within cells from regions of yolk, head and flank of the embryo. We visually inspected each raw cell image with their microridges and eliminated cells with occlusions or very bad contrast on the microridges owed to microscope imaging to maintain high-quality set for the training. However, several pattern configurations with varying but visually discernable local contrasts were considered for the training set. This enabled the network to achieve better learning on the tested hyperparameters required to discern the pattern with variety of complexities. For the convolutional network training, we varied the training set size to ensure the proportions of training set did not largely bias the accuracy values. We randomly divided data into 95%, 93% and 90% as training set and the remaining as test set and tested the same set of hyperparameters on different sized data. The data partitioning was done randomly and for all combinations of tested hyper-parameters and sets. For our segmentation problem, we implemented the encoder-decoder segmentation architecture that performed reasonably well on the dataset.

### Hyperparameter tuning for training

We varied the size of the image, which scales to the receptive field size initially. The original 2D microscope image frames were acquired at 512×512 pixels for all the cells. For the CNN approach, we trained the network with cell image dimensions of 128^2^, 200^2^ and 256^2^. Similarly, we tested the encoder-decoder subnetwork at depths of 2, 4 and 6. We tested the initial learning rate (*ilr*) for 10^−3^ and 10^−4^. Training of all network models was performed using stochastic gradient descent with *ilr*=10^−4^, receptive field size set to 256^2^ and encoder-decoder subnetwork depth at 6. We implemented the median pixel weighted image normalization. A data augmentation approach was included by image rotations, translations and reflections for all training sets and different permutations of hyperparameters. Obtaining the best performance network required empirical analysis and hyper-parameter tuning. For each proportion of the training dataset (90%, 93%, 95%), we varied the mini-batch size (MBS) with values of (6, 9, 11, 15) and number of iterations or maximum epochs (ME) with values of (400, 500, 600, 800) that yielded 16 combinations of these hyperparameters. All training models were run by creating executables using MATLAB compilers to accelerate the training process. The codes were run on a single GPU (NVIDIA Tesla V100 16GB) node on a high-performance cluster. We confirmed the performance of the trained network with reported accuracy using test dataset.

### Segmentation performance evaluation metrics

We implemented the standard evaluation metrics for assessment of image segmentation network performance on test datasets. These included mean pixel accuracy for each class for the entire test dataset, image-wise and class-wise accuracies and Jaccard similarity coefficient (Csurka 2013). The weighted intersection over union (Weighted IOU) measure that weighted the number of pixels for each class was used to benchmark the segmentation performance when comparing results from different combinations of hyperparameters and all proportions of training set. We then evaluated the network performance for MBS and ME for each data proportion.

### Curvature analysis for estimation of microridge persistence length

We describe a method to estimate the distribution of microridge curvature in their dynamical steady state from the skeletonised images of their branches. First, we discuss how to obtain the microridge contour from the images to implement the curvature estimation method. Then we use the experimentally determined curvature distribution to estimate the persistence length and the flexural rigidity, that describes an inherent mechanical property (characteristic length scale and bending rigidity) of the microridges.

### Microridge contour and their orientation

The outline of the microridges was traced to obtain discrete x–y coordinates (*L_xy_*) along the length of a microridge skeleton contour, which is then used to estimate the curvature distribution. To effectively reduce measurement errors in the estimation of curvature, we implemented a ‘cubic smoothing spline’ interpolation method to obtain many smoothened intermediate points on the discrete x–y coordinates obtained from the microridge contour, thereby reducing the spacing between the data points on the microridge trace contour. Also, we ensured that the contours had at the least ten x–y coordinate points prior to implementing the fits in order to reduce errors in the analysis. The interpolation method assumes that the y-coordinate monotonically increases with ‘x-coordinate’ or vice-versa. For inconveniently oriented contours where neither is the case, the method will thus either fail or induce errors in the fit. In order to obtain the correct interpolation on the x-y coordinates, we introduced a 2D rotation matrix *R*(*θ_R_*) for a given angle, *θ_R_*,

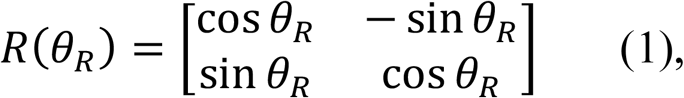

where -π ≤ *θ_R_* ≤ π, and rotated each skeleton coordinate *L_xy_* coordinate in the xy plane by an angle *θ_R_* according to

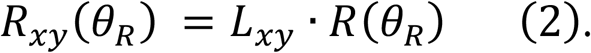

The orientation of the rotated skeleton trace *R_xy_*(*θ_R_*) now depends on the angle *θ_R_*. We varied *θ_R_* over 21 different values and for each implemented the cubic spline interpolation method to obtain a new set of coordinates with several intermediate values given by 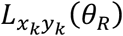. From these, we selected a single 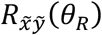 for which the interpolated curve was closest to the original one, thus realising,

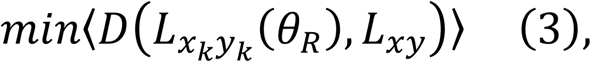

where ‘*D*’ denotes the Euclidean distances between all points in 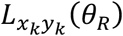 and *L_xy_*. This approach unambiguously provided the best cubic spline interpolation fit on the 2D coordinates extracted from the microridge image data irrespective of their orientation within the 2D images that was then used to estimate the microridge persistence length *(L_p_)*.

### Estimation of persistence length and flexural rigidity from curvature distribution

The classical approaches for the estimation of *L_p_* includes the Fourier shape-fitting method that depends on the number of estimated modes described for actin filaments and microtubules (Gittes 1993) and the curvature distribution method described for DNA chains (Rivetti 1996). For the microridge data, we followed the curvature distribution method, since the Fourier shape-fitting method requires the right number of modes to be fitted, which may not work for all microridge configurations. For a discrete set of two-dimensional coordinates (*x_k_,y_k_*) along the length of the microridges with (*N+1*) points, along the length of the curve (*L*), the spacing Δs_*k*_ between coordinates is given by

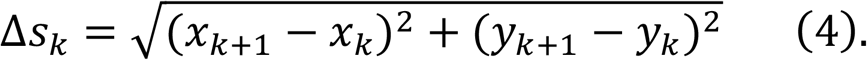

The tangent angle for a set of *N* segments that connect the points is given by

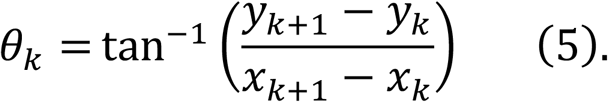

The arc length *L* along the microridge length is given by

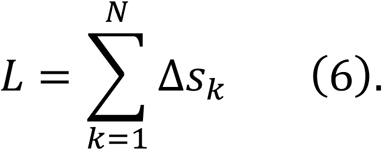

We then estimated *φ*_k_, the angle between two consecutive tangent vectors as

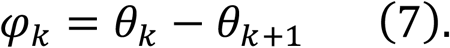

The curvature (*κ*) was approximated for small angle changes and small bond lengths at each coordinate over the average arc length of the two adjacent segments by

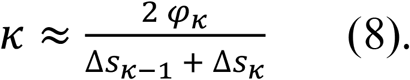

In a subsequent step it will be inconvenient that the segment lengths, Δ*s*_κ_, vary across our ensemble, hence we defined a rescaled variable **κ*_s_*, such that,

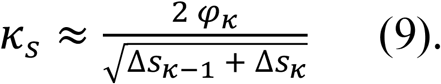

We then defined the effective persistence length (*L_p_*) based on the width of the distribution 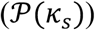 in two dimensions, since the probability of having a certain bending angle follows a normal distribution in thermal equilibrium^39^. The distribution is then given by

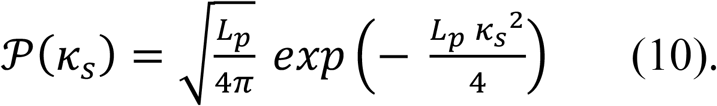

The variance of the distribution is inversely proportional to the persistence length (*L_p_*). The flexural rigidity (*EI*), which characterizes the propensity for thermal bending of a flexible polymer^38^ can then be estimated from

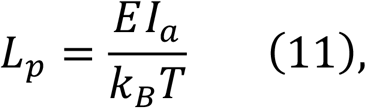

where *k_B_* is the Boltzmann constant, *T* is the absolute temperature in Kelvin (taken as ≈300K, for the imaging temperature of 25°C - 27°C) and the microridge *L_p_* was estimated from the experimental data. *Ε* is actually the Young’s Modulus that relates stress and strain within and *Ι_a_* is the second moment of the cross-sectional area.

### Pattern wavelength

We estimated the overall microridge pattern wavelength for yolk and flank cells in the Fourier domain. For this, we computed the 2D Fourier transform of the network segmented logical images to determine the overall wavelengths of the microridge pattern within each cell. In Fourier space, we calculated a characteristic wave number (*wn*) as described in (Supplementary Methods), from which the corresponding pattern wavelength, *λ*, followed as

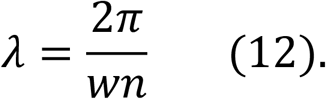

### Branch lengths

The network segmented images were converted to logical arrays and subsequently a ‘skeleton operation’ was used to obtain the traces of the microridges. From the image skeleton, we estimated branch lengths. Branch points were determined and subtracted from the trace images to obtain the skeletonised branches. Branch lengths were computed for each microridge branch by summing up their total number of pixels within a cell. We computed the mean branch lengths considering all the microridge branches for each segmented cell pattern.

### Velocity flow analysis

We estimated the microridge growth velocity (*v_growth_*) and the shrinkage velocity (*v_shrinkage_*). The growth velocity (*v_growth_*) inferred from the image encompassed the merging velocity of adjacent microridges and the local material assembly rates (leading to single microridge elongation events), while the shrinkage velocity (*v_shrinkage_*) accounted for the splitting velocity of microridges and the local disassembly rates (leading to single microridge fragmentation events); all four events are due to net actin dynamics within the microridges. We implemented an optic flow (Lucas-Kanade derivative of Gaussian) method using an inbuilt Matlab function to determine the velocities 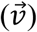 from a time series of spacing *Δt* from network segmented cell patterns with intensity values *I* by computing,

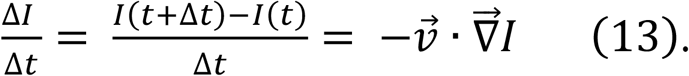

The image velocity divergence 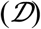 was then computed as

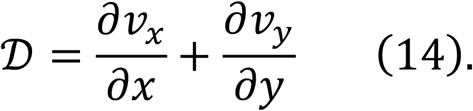

where ∂x and ∂y denote the spatial change in the x and y directions. We implemented a convolution operation using a smoothing kernel σ (where 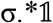, where σ=1/9 and 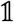 is a 3×3 ones matrix) on the velocity component image matrices for subsequent computations to eliminate low level noise within the image time series. We determined the regions of high positive divergence and strongly negative divergence in the images and determined the velocities specifically at these locations. We then averaged the velocity over all such locations to determine the shrinkage velocities (*v_shrinkage_*) and growth velocities (*v_growth_*) respectively. We computed the strain rate tensor (*S*) considering the symmetric component of the velocity gradient, given by,

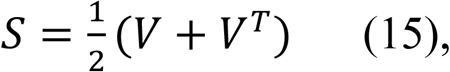

where the velocity gradient (*V*) is,

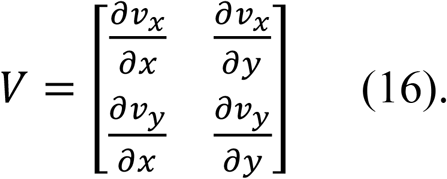

The off-diagonal element of S are the shear strain rates (*γ^S^*) given by,

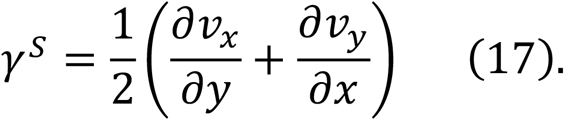

The eigenvalues *⋀_1_* and *⋀_2_* of *S*, such that *⋀_1_>⋀_2_* were computed; *⋀_1_* represent tensile strains, and *⋀_2_* represent compressive strains.

We evaluated the overall principal strains by quantifying the magnitude of cell pattern strains, given by √2||S||, where ||S|| is the matrix norm within microridge regions only, 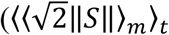 such that 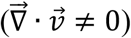 We neglected the anti-symmetric part of the velocity gradient tensor, which represents rotational components, since localized rotational effects were not significant here on their own, as observed in the microridge flow patterns under *wildtype* conditions. The components of S-tensor account for both shape and area changing strain components.

### Area-deviatoric decomposition of the strain rate tensor

The decomposition of the 2D strain rate tensor (*S*) into area strain tensor (*S^area^*) and deviatoric strain tensor (*S^dev^*) can be written as,

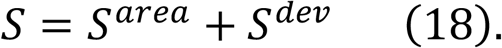

The *S^area^* contains the area changing, shape preserving part, whereas the *S^dev^* contains the shape changing, area preserving part of the total strain rate tensor (*S*).

The deviatoric tensor *S^dev^* is a traceless second order tensor (*tr*(*S*^dev^) = 0) given by,

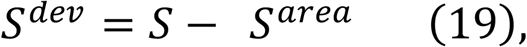

where

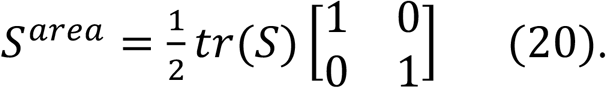

The trace (*tr*) of the strain rate tensor (S) is the image velocity divergence 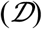 and also the *S^area^* tensor given by,

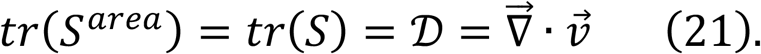

The components of *S^dev^* are purely deviatoric deformations. The eigenvalues of *S^dev^* provide the deviatoric elongation 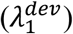 or deviatoric shrinkage 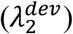 without change in the local area. The eigenvalues of *S^area^* are both equal, and equal to *tr* (*S^area^*)/2.

### Measurement of fluctuations of intensity profiles within microridges

For this analysis, we cropped the center regions of cells within the yolk with relatively shorter microridges to obtain temporal intensity profile within them. We used the detected 2D centroid positions after image segmentation. We wrote custom codes by implementing a multiple hypothesis tracking method frame-by-frame and solved using a linear-assignment problem (LAP) approach with a maximum distance cut-off of ~2.3μm to find the same microridge in the next time frame. Briefly, a cost matrix was computed between centroids of detected microridges that considers all possible assignments between consecutive time frames and assignment is obtained by solving the matrix for minimal cost. A small distance cut-off ensured that tracking errors due to long microridges merging and splitting events were reduced considerably. The purpose of the tracking algorithm implementation was to compute the time evolution of the intensity profile along the microridges. For this computation, the pixel intensity and the eigenvector along the microridge length were used. Generic custom codes were written to obtain the dominant 2D eigenvectors (https://www.mathworks.com/matlabcentral/fileexchange/98894-image2dvectors) of each tracked microridge from the images. For any branched microridge, the intensity profile was assigned along the longer branch, independent of microridge orientation within the image. We obtained the time evolution of the pixel coordinates [*x_p_, y_p_*] and the corresponding intensity readout (*I_pv_*) of microridges on the obtained microridge tracks, and then transformed coordinates into the frame spanned by their eigenvectors. The coordinate along the microridge *δ_l_*, is,

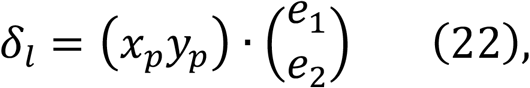

where *e_1_* and *e*_2_ are the components of the normalized eigenvector that points along the length of the microridge. From the mean intensities, we computed an intensity profile along the 1D direction of the microridge, (*I_p_*), by summing the *I_pv_* values for each discretized *δ*_l_.

### Local Gaussian curvature of the microridge pattern on periderm cell surfaces

We discretised the 2D space in (X,Y) and the Z height was taken as the microridge intensity (*I*) readout at each point on the periderm cell surface. We computed the first and second derivatives at each point and the first (*E, F, G*) and second fundamental form (*L, N, M*) of the surface ℝ_3_. Then, the localized Gaussian curvature (*K*) was computed at each point on the cell surface that corresponded to the localized intensity within the microridges, given by,

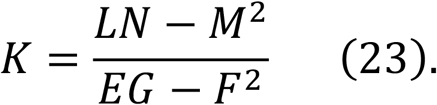

Gaussian curvature (https://www.mathworks.com/matlabcentral/fileexchange/11168-surface-curvature) was modified to compute the Gauss gradient with σ=1.2μm using (https://www.mathworks.com/matlabcentral/fileexchange/8060-gradient-using-first-order-derivative-of-gaussian) to extract the first and second derivatives at each point in the image.

### Quantitative parameter analyses of microridges from network segmented images

The CNN approach for training and test data consisted a total of 1502 single periderm cell images patterned with microridges. Using the best performing trained network, we segmented microridge of image sizes 256^2^ and the physical size of image pixels were rescaled for each image prior to parameter extraction (Supplementary Methods). We considered 593 cells (300 yolk and 293 flank cells) comprising of thousands of microridges for the estimation of persistence lengths (*L_p_*) of microridges. The number of microridges in this analysis was kept large for estimation of bending rigidity from the experimental data. The comparative analysis of static parameters of microridges included the overall pattern wavelength (*λ*) and mean branch lengths (*<B_l_>*) on 300 yolk cells and 293 flank cells. For the steady state dynamic parameter analysis, we used the time series data from yolk and flank regions. We examined 9 movies with tracks of varying length durations between 11 minutes and 32 minutes. All the computational analysis was performed in Matlab (Mathworks) using custom codes.

## Supporting information

BhavnaR_SonawaneM_Supplementary_Movies

## ACKNOWLEDGMENTS

Bhavna R is thankful to Yogesh Arora, Bhatkar PJ and their team of the TIFR Central Workshop for fabrication of the mounting piece for imaging zebrafish embryos, Anil Kumar Naik and the TIFR Computer Centre group for their help and support in providing GPU and HPC cluster for CNN training. We acknowledge Boby KV for taking care of the light microscopy, Kalidas Kohale for fish facility maintenance and the DBS kitchen staff for common reagents. We thank current and past members of the Sonawane laboratory for discussions. We acknowledge the department of atomic energy (DAE), TIFR for the funding (RTI4003;12P-121). We thank Mandar Inamdar for insightful discussions and valuable comments on the manuscript. We thank Sebastian Wüster for valuable comments and critically reading the manuscript.

## AUTHOR CONTRIBUTIONS

RB- Conceptualization, methodology, investigation, data curation, formal analyses, visualization, writing manuscript draft and editing; MS- laboratory support for experiments, draft editing, revision.

## Supplementary Information

### SUPPLEMENTARY METHODS

We describe the details of the image processing pipeline to produce a segmentation mask of the microridges. The algorithm consists of a 2-step segmentation approach. In the first, we segmented the periderm cells, followed by cell tracking and frame by frame linkage. This was then followed by a tailored segmentation algorithm to label the intracellular pattern of microridges within each cell, enabling a rapid annotation process. The single periderm cell extraction patterned with microridges formed the training datasets for convolutional neural network segmentation.

#### 1. Image processing pipeline for microridge segmentation

##### 1.1 Peridermal cell image filtering

The images were obtained from periderm cells up to the basal epidermis in the z-direction in XYZT formats (Supplementary Fig S1a). In order to filter out the periderm slices from the basal epidermis, we implemented a 2-step image entropy-based filtering.

The filtering parameter were adjusted separately for cells of yolk and flank images empirically for each confocal image series because the movement along the z-direction (because of tissue thinning along the z-axis) was different within different regions of the embryo and varied over time. Additionally, the yolk cells are relatively taller than the flank cells and this affected how quickly we reached from periderm to basal epidermis in the z-direction. First, the global entropy for each slice was computed using the Shannon information content (Matlab function, *entropy*), since the slices of periderm cells have relatively lower entropy than the z-slices from the basal epidermis. Therefore, an entropy threshold after manual inspection was set to filter out periderm slices from the basal epidermis. Secondly, in some regions of the same z-slice (middle slices) there was overlap between periderm of one cell and basal epidermis of another cell within the same slice of the flank region. In order to filter out distortions from the basal regions, the local entropy within a 3×3 window size (using Matlab function, *entropyfilt*) for the selected slices from the previous step was computed. This step was not that crucial for the yolk cell patterns because of their relatively greater cell heights from the basal epidermis. A pixel-level entropy threshold was then used to filter out noisy pixels of the basal cells that corrupted the microridges and periderm cell membrane. The noisy pixels coming from the basal layer typically were locally of higher entropy. Both these parameters were set after manual inspection for each confocal image series, and kept constant throughout all time points, to extract periderm cells only (Supplementary Fig S1b-c).

##### 1.2 Periderm cell segmentation

The mean intensity along the z-direction of filtered periderm slices was computed and the data for image segmentation was reduced to XYT dimensions (Supplementary Fig S1d). We used a low pass, linear, Gaussian smoothing filter given by,

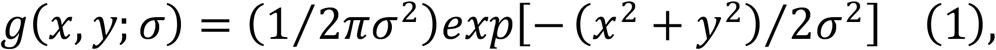

with a standard deviation of σ=0.7 pixels to reduce noise in all the images.

Next, to demarcate the cell membrane boundary only, the microridges were eliminated (momentarily) from within the cells. For this, we applied a Butterworth high pass (BHP) frequency filter of order *n* and cut-off frequency D_0_ defined as

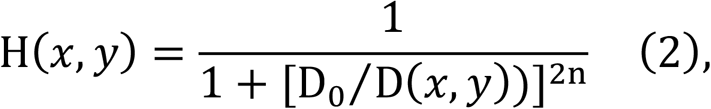

where *D(x,y)* is given by,

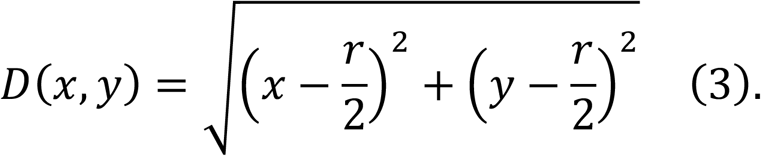

Here, *r, c* are the image width and height in pixels, respectively (Supplementary Fig S1e). The filter order, *n* =1 and D_0_=3 (https://www.mathworks.com/matlabcentral/fileexchange/40579-frequency-domain-filtering-for-grayscale-images). This step was followed by image binarization and morphological operations to close any gaps on the cell membrane due to varying image contrasts. The parameters for morphological operators, *imclose* and *strel* Matlab functions were adjusted depending on the image contrast, and then kept constant throughout all time points for a single movie. Each enclosed region (cell) bounded by the cell membrane was masked using the complementary image (*imcomplement* function) as shown in (Supplementary Fig S1f).

##### 1.3 Single periderm cell extraction

The *Area* and *Solidity* properties for each cell enclosed by completed boundaries were computed. We implemented suitable cut-off values on the ‘cell area’ and ‘solidity’ to eliminate cells only partially visible at imaging borders. The cells for which the cell boundaries were incomplete were automatically discarded in this process. For the remaining selected cells, enclosed by completed boundaries, the ‘BoundingBox’, ‘ConvexImage’ and ‘Centroid’ properties were computed. The *BoundingBox* finds rectangular coordinates for each enclosed cell region. Each ‘BoundingBox’ coordinate corresponded to a single cell and was used to extract rectangular regions of a cell using the *imcrop* function on the original Gaussian smoothened image comprising of both microridges and cell membrane. This allowed the isolation of each cell with their microridges from the initial images as shown in Supplementary Fig S1g. Cell shapes are typically polygonal, however the bounding box coordinates resulted in rectangular selection around each cell. In order to mask the cell enclosed by their membrane, the *ConvexImage* was used. The *ConvexImage* is a binary image with only enclosed pixels of a cell bounded by its membrane set to 1’s (shown in yellow). The Hadamard matrix product of cropped rectangular cell region and its *ConvexImage* resulted in selective extraction of individual periderm cells over time (Supplementary Fig S1g).

##### 1.4 Periderm cell tracking

In order to track the same periderm cell frame to frame, a nearest neighbor approach based on the Euclidean distance between cell centroids at time *t*_*n*_ and *t*_*n*+1_ was implemented. The cell tracking allowed us to follow the same cell patterned with their respective microridges as obtained from the microscope (Supplementary Fig S1h).

##### 1.5 Microridge segmentation: labeled images for the training set

For each extracted periderm cell, we used a Gaussian smoothing filter (σ=0.7 pixels) given by Equation (1) to smoothen the microridges pixels and filter out low background noise from raw cell patterned images (*R*). We convolved the resultant Gaussian smoothened image, *IM*(*x*, *y*) with the derivative of Gaussian ∂*g*(*x*,*y*;*σ_g_*) to obtain the microridge intensity gradient,

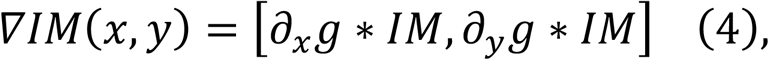

where * denotes convolution (Supplementary Fig S1g).

We computed the second derivative of the Gauss gradient image to obtain the Hessian matrix at each pixel. The trace of the matrix gives the image Laplacian given by,

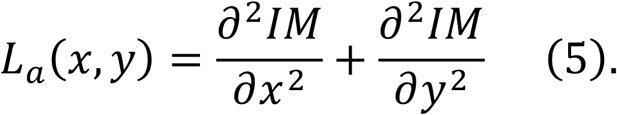

We considered only negative Laplacian values *L(x,y)*,

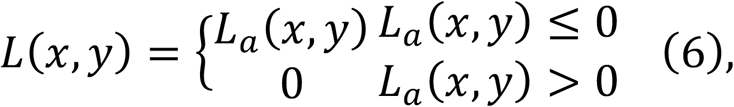

in order to select regions with negative intensity emphasizing only on microridges. Subsequently, in order to obtain a smoothened image segmentation mask, we fitted the *L(x,y)* matrix into a logistic sigmoid function given by,

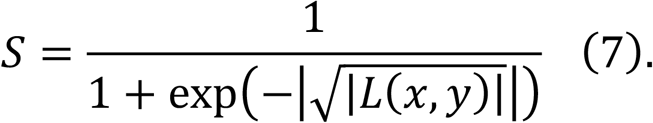

As a next step, we generated a binary image using the *imbinarize* Matlab function on the image *S*. This resulted in a labeled binary image (*B*) of microridges (Supplementary Fig S1i). The image processing pipeline described so far was used to create a labeled training dataset for neural network pixel-wise semantic segmentation. Pairs of raw extracted cells patterned with microridges and their corresponding labeled image (*B*) formed the training set for the CNN based microridge segmentation framework.

#### 2. Extraction of quantitative parameters for segmented microridges

##### 2.1 Generation of network segmented cells patterned with microridges

After successful training, the networks generated segmented periderm cells patterned with microridges in the form of binarized images (*NB*). We created masked cell image (*NM*) given by the Hadamard matrix product of raw cell patterns (*R*) and the corresponding network produced image binary (*NB*),

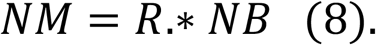

The masked image (*NM*) is defined on only those gray pixels in the original image where the *NB* image takes values of 1.

##### 2.2 Adjustment of pixel sizes of network segmented images

All the network segmented images (*NM* or *NB*) were scaled to dimensions 256×256 in xy in order to obtain high performance on the network training. The original microscopy movies were acquired at spatial dimensions 512×512 and consequently, original cell sizes were of smaller dimensions. Hence, prior to computing quantitative parameters of microridges, we calculated the physical pixel size in the x and y directions from the original cell dimensions. All confocal images were acquired at pixel size, *psz*=0.1977μm in both xy directions. Hence, for an extracted cell image dimension (*x*_o_, *y*_o_), we re-scaled the pixel sizes Δ***x*** and Δ*y* in microns in x and y according to new image sizes of 256×256, given by,

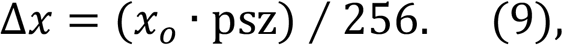

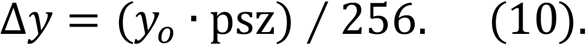

##### 2.2 Fourier transform approach for determining pattern wave number

The 2D Fourier transform of the image *I*(*x*, *y*) was calculated using the Matlab *fft2* function, to compute,

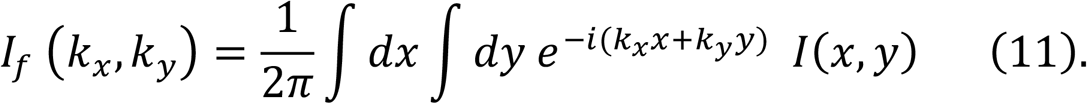

The Fourier space of *k_x_* and *k_y_* ranges from −K to K, where 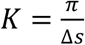, and Δ*s* is the edge lengths of 1 pixel within the image. We defined the characteristic pattern wave number (*wn*) by,

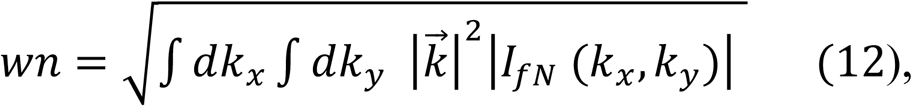

where,

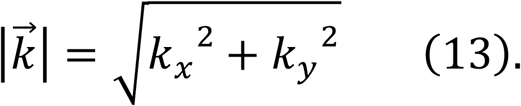

The magnitude of the image Fourier transform was normalized given by,

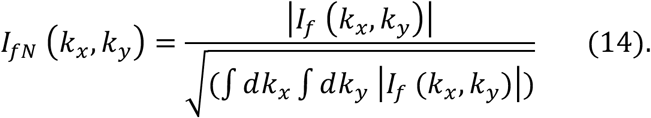

### SUPPLEMENTARY FIGURES

**Fig S1.**
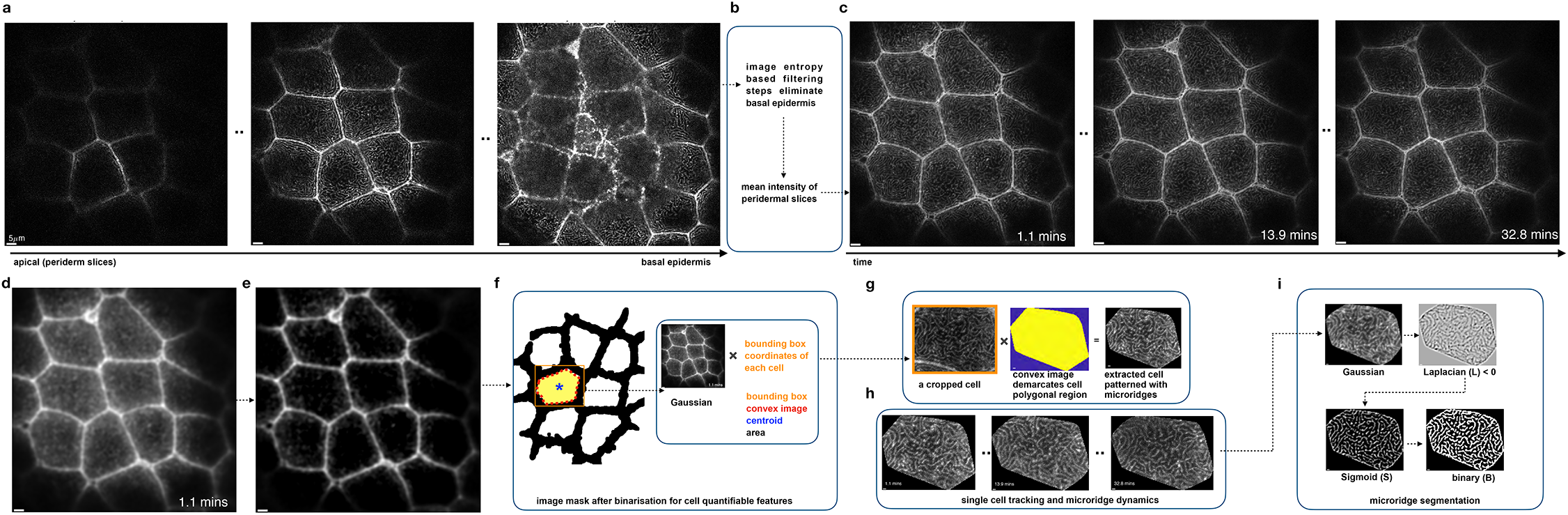
Microridge image processing pipeline forms the training set for the CNN approach. **a**. Live imaging of zebrafish embryonic epidermis at 48hpf was performed to acquire images of periderm cells until 4-5μm depth reaching the basal epidermis. **b**. Gaussian image slices (σ=0.5 pixels) were processed using global entropy filtering to eliminate basal epidermis slices. For tackling periderm and basal signal on the same slice, a local entropy filtering threshold was used to eliminate the noisy signal from the basal slices. **c**. Time-lapse images of mean intensity periderm slice images **d**. Gaussian convolved (σ=2.5 pixels) filtered periderm cells **e**. Processed images using a high pass frequency filter that eliminated only microridges pixels while preserving the cell membranes. **f**. Image binarization demarcated cell membrane boundary (red dotted line) and the following cell features were extracted: bounding box rectangular coordinates (orange); convex cell image (filled-in yellow); cell centroid (blue asterisk) and cell convex area. Product of bounding box rectangular coordinate of each segmented cell and the Gaussian image (in **b**.) extracted each cell within the box coordinates from the time-lapse images. **g**. Hadamard product of rectangular demarcated cell (with microridge) and each cell convex image extracted single cell images with their microridges that formed the raw input training set for the CNN approach. **h**. Centroid based cell tracking was implemented to follow the same cell over time and their microridge pattern dynamics. **i**. Microridge segmentation was achieved by employing sequential steps: Gaussian, followed by negative Laplacian image, processed further using a sigmoid function. Image binary was obtained from the sigmoid image that formed the labeled training set for the CNN approach. Details are given in Supplementary Methods.

**Fig S2.**
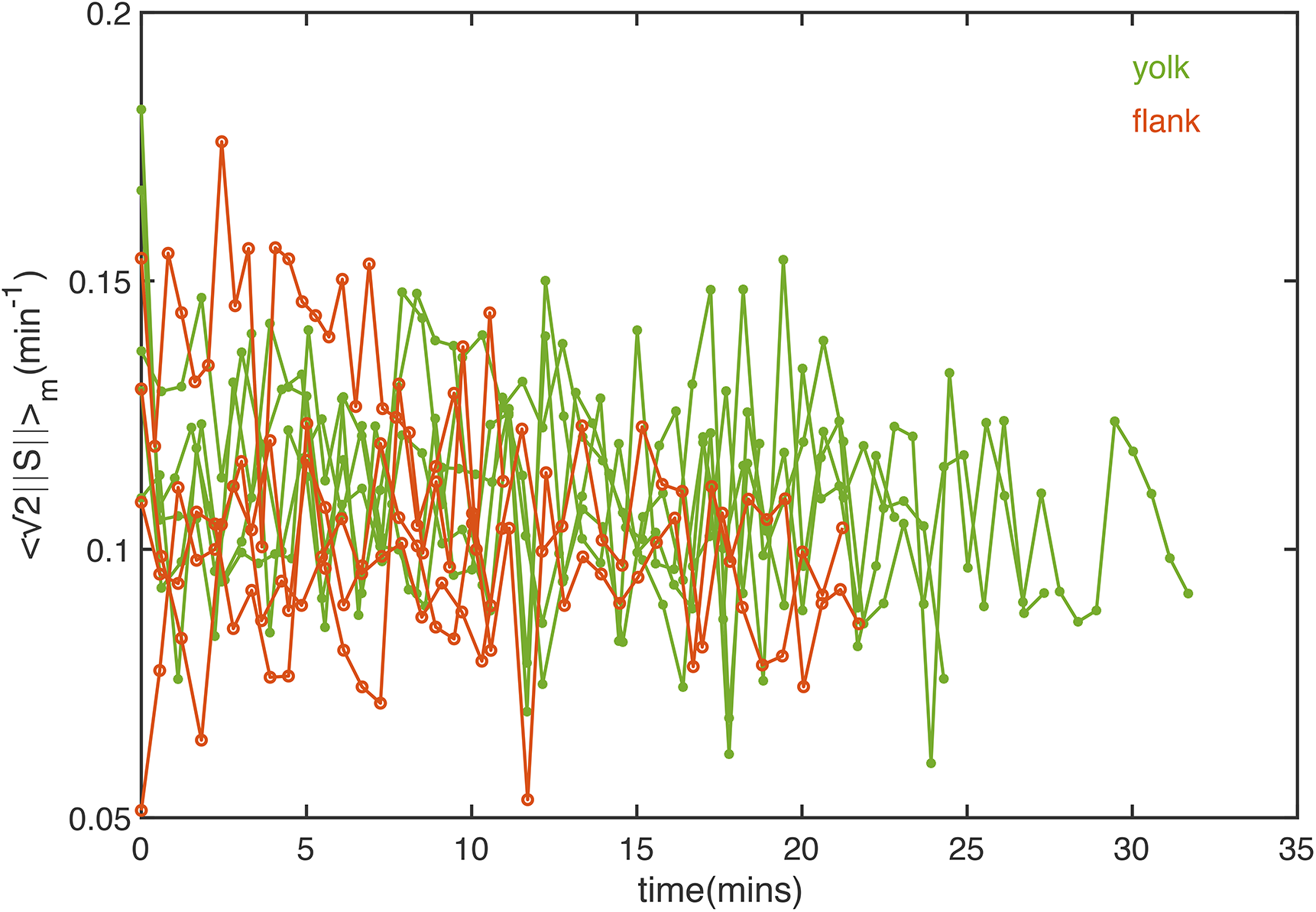
Strain rate tensor norm of microridge cells patterns from yolk versus flank regions. Temporal evolution of the norm of the strain rate tensor averaged over microridge patterns, given by (√2||S_*m*_||), (where *m* stands for microridge regions only) computed for both yolk (green) and flank (orange) cell patterns. Generally, strain-related parameters of the yolk cell patterns are larger than in patterns from the flank regions. In both cases, the mechanical parameters of cell surface patterns revealed the presence of temporal periodic fluctuations indicating mechanical oscillations.

**Fig S3.**
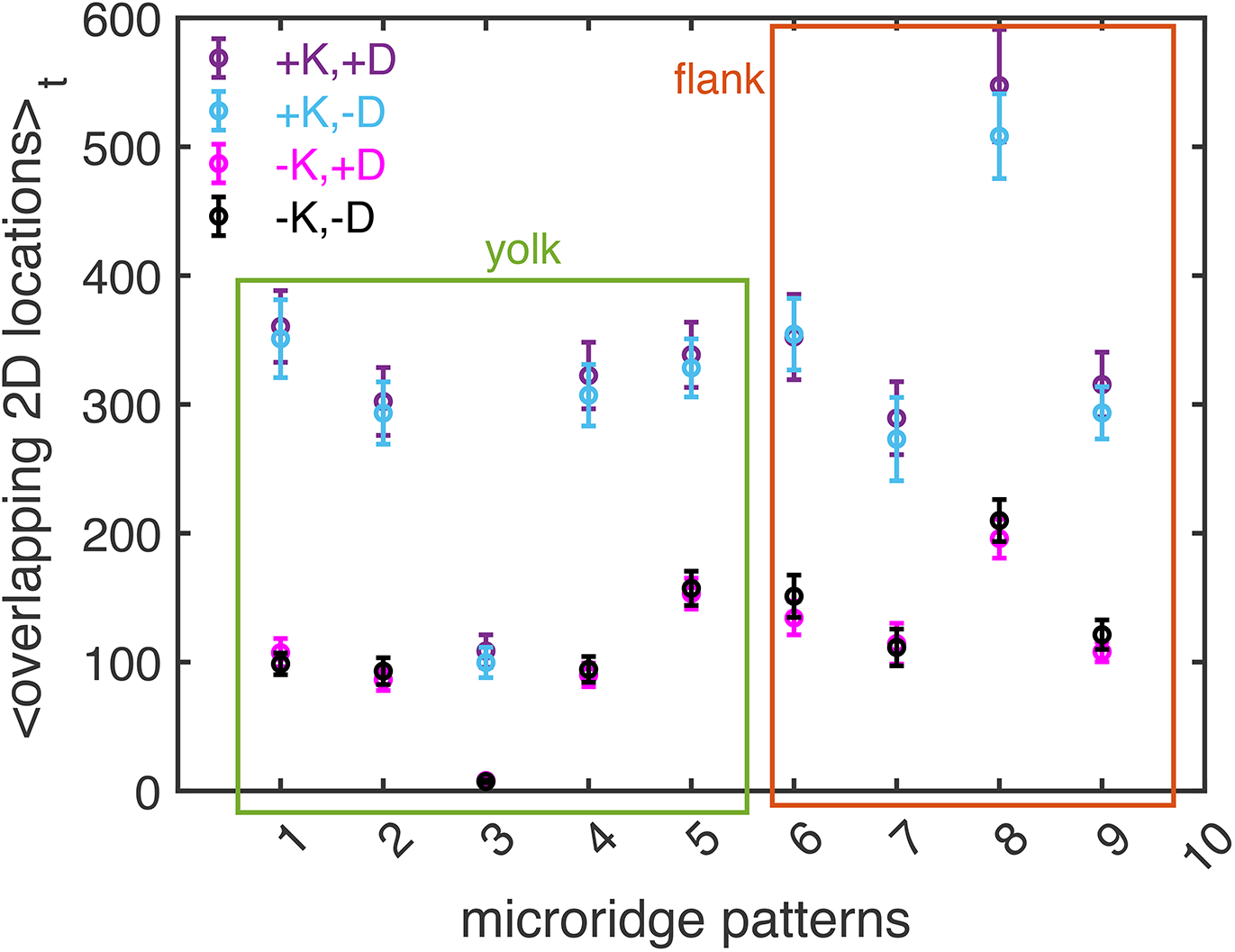
2D spatial co-occurrences of localized Gaussian curvatures (K) and the Divergence 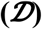 of the cell pattern dynamics. Each data point indicates the mean number of coincident 2D-locations and their standard error of mean for a cell pattern dynamics from yolk or flank region. In all cases, higher frequency counts of +*K* locations at time *t* overlapped with ±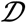 locations from time *t* to (*t+1*).

### SUPPLEMENTARY TABLE

**Table S1.**
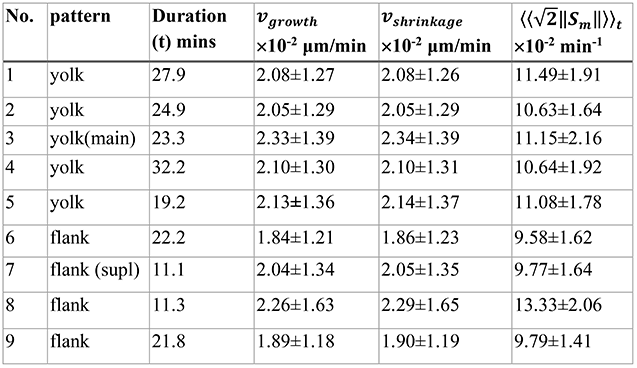
Velocity flow analysis parameters for yolk and flank cell microridge patterns. The parameters *v_growth_*, *v*_shrinkage_ and (√2||S_*m*_||) were computed for 9 cell patterns. The overall strain rates built-up within the patterns of yolk are relatively higher than flank cell pattern, as indicated by the computed parameters.

### SUPPLEMENTARY MOVIES

**Supplementary Movie 1**. A movie of a tracked yolk cell patterned with microridges and the corresponding convolutional neural network segmented movie.

**Supplementary Movie 2**. A movie of a tracked flank cell patterned with microridges and the corresponding convolutional neural network segmented movie.

**Supplementary Movie 3a**. A cropped square region of the yolk cell (shown in Supplementary Movie 1) center illustrating microridge pattern dynamics.

**Supplementary Movie 3b**. Panel 1 shows a cropped square region within the yolk cell (shown in Movie 3a) center illustrating microridge pattern dynamics at time *t* and overlay of the velocity vectors at time (*t+1*) in magenta. Panel 2 is the divergence 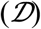 given by the trace of strain rate tensor (S). Panels 3-4 correspond to time series analysis of the tensile strains 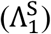 and compressive strains 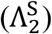 of microridges within the yolk cell pattern. Panels 5-6 give the deviatoric elongation strain 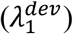 and deviatoric shrinkage strain 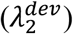 without change in local area. The colorbar axis range was set for visualization across all time frames.

**Supplementary Movie 4a**. A cropped square region of the flank cell (shown in Supplementary Movie 2) center illustrating microridge pattern dynamics.

**Supplementary Movie 4b**. Panel 1 shows a cropped square region within the flank cell (shown in Movie 4a) center illustrating microridge pattern dynamics at time *t* and overlay of the velocity vectors at time (*t+1*) in magenta. Panel 2 is the divergence 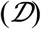 given by the trace of strain rate tensor (S). Panels 3-4 correspond to time series analysis of the tensile strains 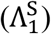 and compressive strains 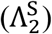 of microridges within the flank cell pattern. Panels 5-6 give the deviatoric elongation strain 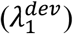 and deviatoric shrinkage strain 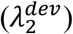 without change in local area. The colorbar axis range was set for visualization across all time frames.

**Supplementary Movie 5**. Intensity readout along a microridge length exhibits positional fluctuations of high intensity of accumulated actin clusters.

**Supplementary Movie 6**. Left panel shows the microridge pattern dynamics within a sub-region of a yolk cell pattern and the right panel are the corresponding localized *K*-values, highlighting regions of high and low intensity within the microridges. The z-intensity profile is shown between 0.2 and 1 to clearly show the high intensity regions.

